# From miRNA target gene network to miRNA function: miR-375 might regulate apoptosis and actin dynamics in the heart muscle via Rho-GTPases-dependent pathways

**DOI:** 10.1101/2020.10.20.344556

**Authors:** German Osmak, Ivan Kiselev, Natalia Baulina, Olga Favorova

## Abstract

MicroRNAs (miRNAs) are short single-stranded non-coding RNA molecules, which are involved in regulation of main biological processes, such as apoptosis, cell proliferation and differentiation, through sequence-specific interaction with target mRNAs. In this study we propose a workflow for predicting miRNAs function by analyzing the structure of the network of their target genes. This workflow was applied to study the functional role of miR-375 in the heart muscle (myocardium), since this miRNA was previously shown to be associated with heart diseases and data on its function in myocardium are mostly unclear. We identified *PIK3CA, RHOA, MAPK3, PAFAH1B1, CTNNB1, MYC, PRKCA, ERBB2*, and *CDC42* as key genes in the miR-375 regulated network and predicted the possible function of miR-375 in the heart muscle, consisting mainly in the regulation of the Rho-GTPases-dependent signalling pathways.

We implemented our algorithm for miRNA function prediction into Python module, which is available at GitHub (https://github.com/GJOsmak/miRNET)

## Introduction

MicroRNAs (miRNAs) are a class of short single-stranded non-coding RNA molecules about 21-25 nucleotides in length, which regulate gene expression at the post-transcriptional level through sequence-specific interaction with mRNAs of target genes followed by their degradation or repression of translation [1]. MiRNAs are involved in regulation of main biological processes, such as apoptosis, cell proliferation and differentiation [2]. In the cardiovascular system, miRNAs play essential roles in the growth and development of cardiomyocytes, contractility, heart rhythm control, angiogenesis and metabolism of lipids [2–4].

We previously found the association of circulating miR-375 level with myocardial infarction (MI) [5]. The study [6] showed the possible functional role of this miRNA in the heart muscle through the regulation of the PI3K/Akt signaling pathway involved in many biological processes, in particular, cell proliferation and differentiation. Other than that, data on the miR-375 function in myocardium are lacking.

In this work, we investigate the possible function of miR-375 in the heart muscle by analyzing the structure of the network of tissue-specific target gene interactions.

## Methods

### Databases

Human Protein Atlas database was used to extract data on gene expression of detected protein coding genes in heart muscle [7]. MiRTarBase database [8] was used to select direct target genes of miR-375. STRING database [9] was used to find molecular interactions between protein products of these genes.

### Network construction and analysing

Gene-gene interaction network was constructed using the NetworkX 2.0 package for Python [10]. Target genes served as nodes of the constructed networks, whereas molecular interactions of their protein products served as edges. Splice variants of one gene were combined into the one node. Since direct targets of miRNAs frequently behave as network hubs or hubs-bottlenecks [11], centrality of nodes was calculated as an additive sum of their betweenness and degree centralities; it was implemented through betweenness_centrality() and degree_centrality() functions in Python package NetworkX. The resulting networks and their characteristics were visualized using Cytoscape software [12].

### Key nodes selection

For key nodes selection we ordered all of nodes by decreasing of their centrality and construct function

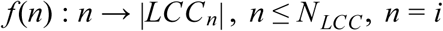

where: *n* - number of top centrality nodes removed,

*i* - index of nodes

*N*_*LCC*_ - number of nodes in largest connected component (LCC)

|*LCC*_*n*_| - cardinality of the LCC from network with removed *n* top centrality nodes

Next, we found the minimal *n* when derivative of this function is equal to zero:

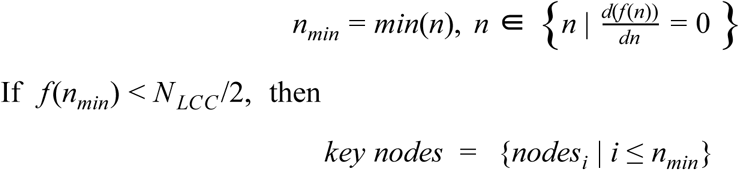

If *f*(*n*_*min*_) < *N*_*LCC*_/2, then

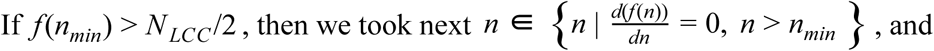

key nodes is a subset of a set of centrally ordered nodes such that the index of the nodes of this subset does not exceed the *n*_*min*_.

If *f*(*n*_*min*_) > *N*_*LCC*_/2, then we took next 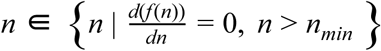, and repeated the procedure.

### Reactome gene set enrichment analysis

For gene set enrichment analysis we used reactome API, which was implemented in the Reactome2py v.1.0.1 package for Python3 [13]. For visualization of the results we used ReacFoam, high level pathways overview visualisation based on Voronoi tessellation, and implemented in Reactome tools.

Gene set clusterization and dendrogram visualization were performed based on the method described in [14].

### Availability of data and materials

The code is available at GitHub: https://github.com/GJOsmak/miRNET.

## Results

Using the Human Protein Atlas database we selected a total of 7944 protein-coding genes, expressed in heart muscle. In this tissue according to the MiRTarBase database, there were 267 target genes of miR-375 (about 3.5% of the total number of protein-coding genes). To evaluate the possible functional role of miR-375 in the heart muscle, we analyzed the structure of the gene-gene interaction network, composed of these 267 genes. Of them, 82 genes are indexed in the String database and have at least one connection. The largest connected component (LCC) of the network contains 53 genes (Figure 1A). Other connected components contained less than 6 genes each. If extracting random gene sets equal in cardinality to the considered sets of miR-375 target genes, the probability of getting LCC of the same sizes is less than 0.0001 that corresponds to the level of statistical significance *p* <0.05.

**Figure 1.**
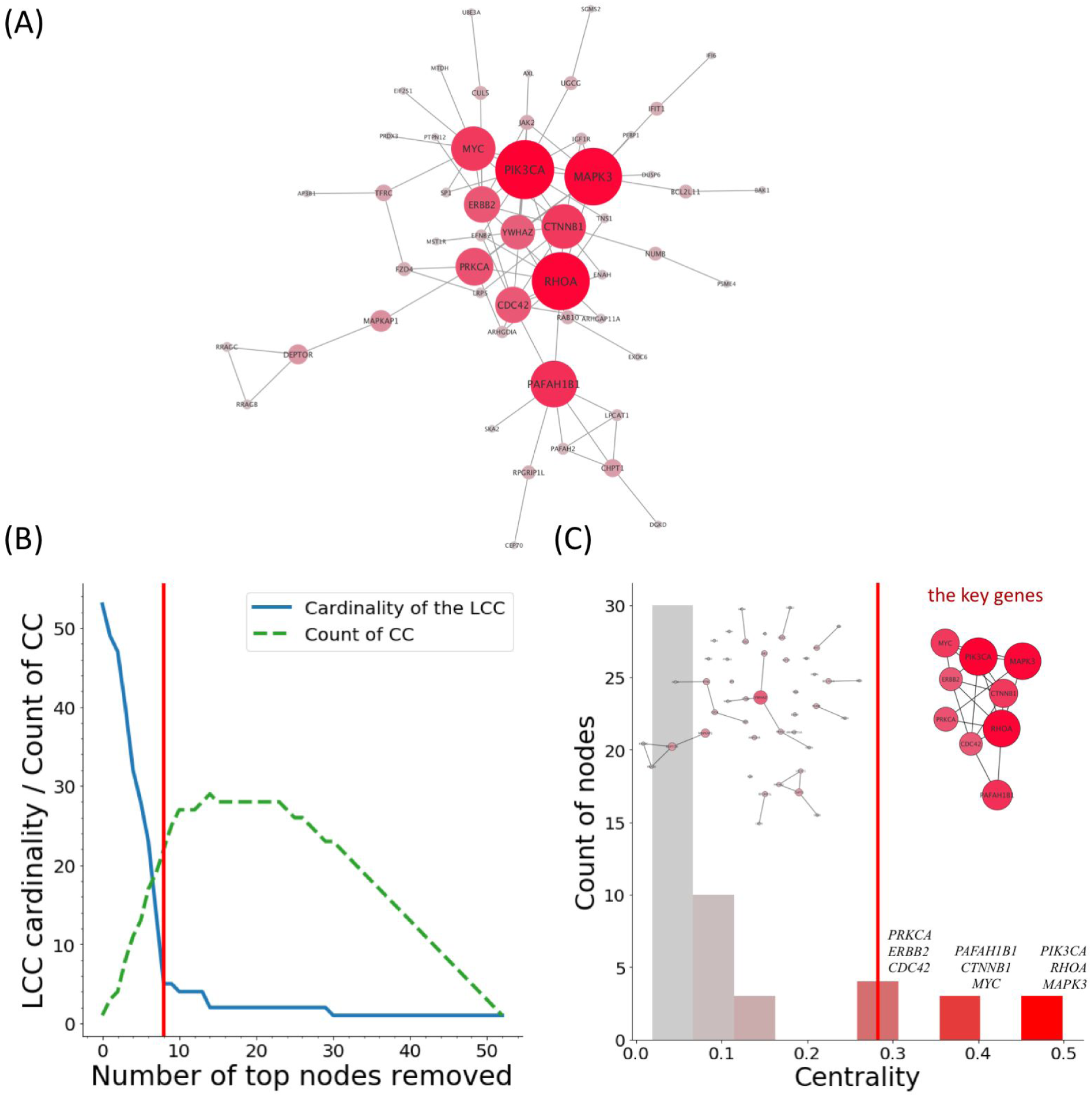
Selection of key protein-coding miR-375 target genes expressed in the heart muscle. The largest connected component (LCC) of the network of miR-375 protein-coding target genes (A). After one by one removal of the genes of the highest centrality, the LCC disintegrates and loses its connectivity. The plot (B) shows the reduction of the LCC cardinality (blue line) and simultaneous enlargement of the connected components (CC) number (green dashed line) during nodes’ removal. The histogram (C) of centrality distribution of genes in the LCC; top genes essential for connectivity and corresponding parts of the LCC: all but key nodes (left network) and key nodes only (right network). The vertical red lines on figures (B) and (C) mark the cut-off level for selection of the key genes when the LCC cardinality hits a plateau. Differences in color of nodes (A, C) and columns (C) from gray to red mark the increase in centrality of genes.

To identify key genes of miR-375 regulated network we were extracting top genes one by one by the centrality of their nodes from the LCC until the LCC cardinality hit a plateau (Figure 1B). Extracted key miR-375 target genes, essential for the connectivity of the LCC (in descending order of centrality), are *PIK3CA, RHOA, MAPK3, PAFAH1B1, CTNNB1, MYC, PRKCA, ERBB2, CDC42* (Figure 1C).

To investigate main functions of the key miR-375 target genes, we performed their overrepresentation analysis in Reactome signaling pathway gene sets. In total, the key genes are significantly overrepresented (*p*-value <0.05) in 94 signaling pathways (Figure 2).

**Figure 2.**
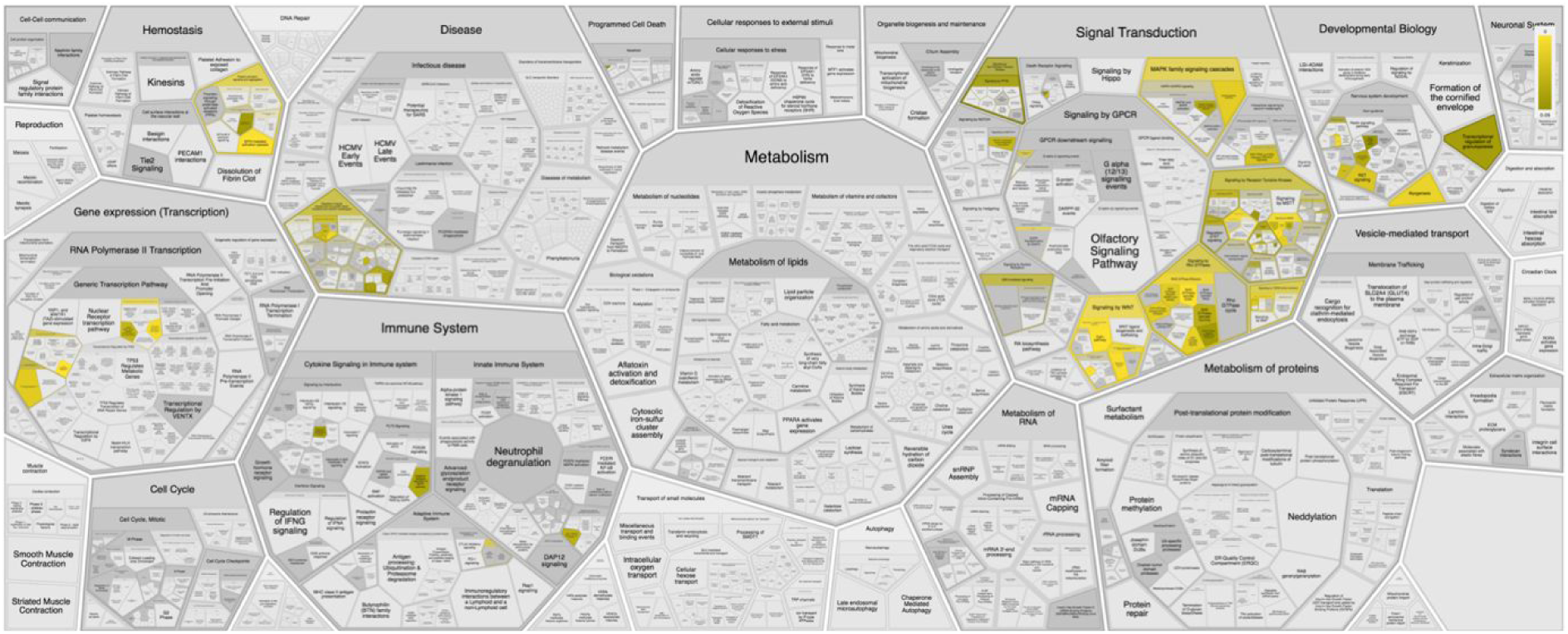
ReacFoam of cell signaling pathways, in which 9 key miR-375 target genes of high centrality are overrepresented. The p-values are shown with the gradient from gray (*p* ≥ 0.05) through brown to yellow (*p* → 0).

As seen from Figure 2, the considered genes are most involved in various signal transduction pathways. Many of the genes of high centrality encode transcription factors located at the very end of signal transduction pathways bearing the corresponding names, particularly RHO-, MAPK-, PI3K/AKT-, MYC- and ERBB2-signaling pathways.

Further, we ranked the overrepresented reactome pathways by the number of key miR-375 target genes and filtered out those that include less than three of the key genes. It allowed us to exclude from consideration side pathways and to focus only on pathways that are to a greater extent represented by key miR-375 target genes (Figure 3).

**Figure 3.**
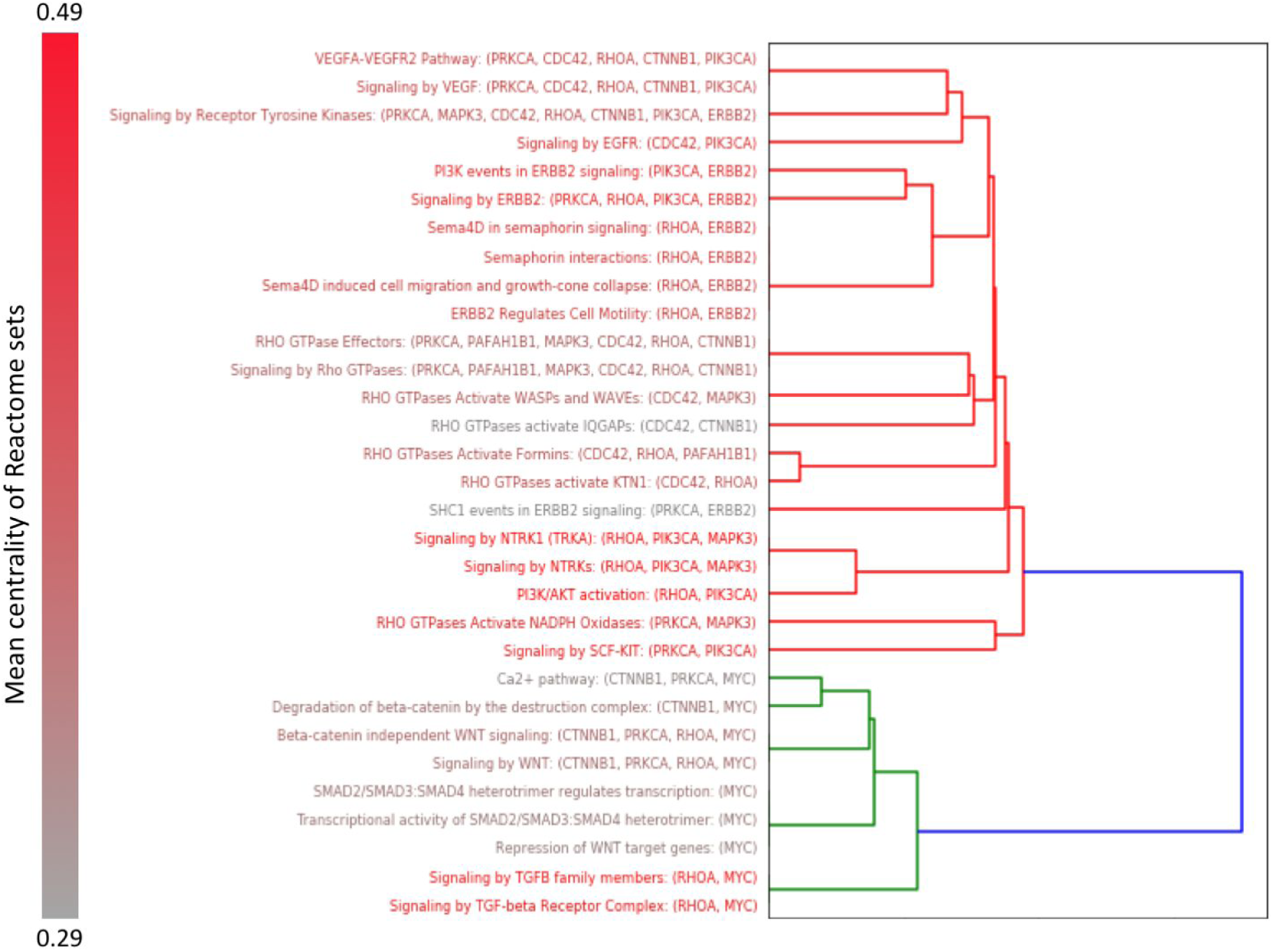
The dendrogram shows clusterization of Reactome pathways, in which the key miR-375 target genes are significantly overrepresented in the heart muscle. The clusters are shown in different colors. Red to gray gradient of text color (shown on the left) reflects the increase in the mean centrality of genes in sets.

As can be seen from Figure 3, signaling pathways containing key miR-375 target genes are divided into two large clusters. The first cluster mainly includes the Rho-, NTRK-, ERBB2-, PI3K/AKT-signaling pathways (red branches of the dendrogram); the second - TGFb-, WNT- and SMAD-signaling pathways (green branches of the dendrogram). All reactome sets from the second cluster include *MYC* gene, and sets with high mean centrality among them also include *RHOA* gene. In turn, all sets from the first cluster, except for EGFR- and ERBB2-signaling pathways, include at least one gene from the Rho kinase signaling pathway (*RHOA* or *CDC42*); many sets with high mean centrality from the first cluster also include *PIK3CA* gene. It should be noted that PI3K/AKT- and MAPK- pathways are located lower in the signal transduction cascade than Rho-GTPases pathways, and form intersection points of the latter with other signaling pathways. Thus, most of the signaling pathways from both clusters of the miR-375 key target genes are associated with Rho-GTPases-dependent signal transduction cascades.

## Discussion

In this study we analyzed gene-gene interaction networks of miR-375 target genes expressed in the heart muscle based on data of gene expression of protein coding genes in this tissue from the Human Protein Atlas database. Results of the statistical overrepresentation analysis demonstrate that key target genes of miR-375 are involved in intracellular signal transduction, predominantly through Rho-GTPase-signaling. PI3K/AKT-, MAPK-, WNT-, NTRK-, ERBB2-, and EGFR- signaling pathways were also overrepresented with key miR-375 target genes. However, PI3K/AKT-, MAPK- pathways are located downstream in the Rho-GTPase signal transduction cascade, whereas WNT-, NTRK-, ERBB2-, and EGFR-pathways may function via Rho-GTPase signaling transducers [15–19]. Thus, we may conclude that miR-375 action in the heart muscle is primarily directed to the regulation of Rho-GTPase genes’ expression.

Rho-GTPases represent a distinct family within the superfamily of Ras-related small GTPases. This family includes small (about 21 kDa) proteins that function as binary switches involved in regulation of main cellular processes such as morphogenesis, polarity, movement, and cell division [20]. *CDC42* and *RHOA* encode proteins that along with Rac1 are the most studied members of the Rho-GTPase family. They induce rearrangements of the actin cytoskeleton and regulate the contractility of actomyosin and the dynamics of microtubules [21,22].

The role of small GTPases in the development of cardiovascular diseases is described [23]. In model animals with ischemia-reperfusion injury inhibitors of RhoA/ROCK signaling pathway demonstrate cardioprotective effect, which is realized through activation of MAPK and NFkB signaling pathways and reduction of Bcl-2 levels leading to a decrease in apoptotic activity of cells [24–28].

In this work, analyzing the structure of the network of miR-375 target genes we showed that its function in the heart muscle may be mainly to regulate Rho-GTPases-dependent signaling pathways. Indeed, *CDC42* and *RHOA* genes which are the direct targets of miR-375 occupy central positions in interaction networks (see Figure 1). Our data are in good agreement with the recent data on the protective effect of miR-375 overexpression in cardiomyocytes after hypoxic-reoxygenation injury [29].

Along with the cardioprotective effect of miR-375 overexpression, its downregulation also showed protective effect but in another experimental model: in mice with MI caused by ligation of the left anterior descending artery without reperfusion [6]. This contradiction might be explained based on our data taking into account a broad variety of molecular functions of Rho-GTPasesin the cell. Probably, inhibition by miR-375 of those Rho-GTPases which are apoptosis activators [30] may have the positive effect in ischemia-reperfusion injury of cultured myoblasts, previously demonstrated in [29]. At the same time, miR-375-dependent inhibition of GTPase Cdc42, one of the key intracellular messengers in angiogenesis [31], may have the negative effect on MI recovery, earlier observed in model mices [6].

In general, this study shows a workflow for predicting miRNA function in different tissues and cells by analyzing the network structure of its target genes. This approach allowed us to predict the possible function of miR-375 in the heart muscle, realized via the regulation of the expression of Rho-GTPase effectors.

## Limitations and future direction

The predicted function of miRNA-375 certainly needs experimental verification. The use of binary estimates of miRNA target gene expression levels in tissue could lead to the loss of some information.

## Abbreviations

MI: Myocardial Infarction
LCC: The Largest Connected Component
CC: Connected Components
miRNA: microRNA

## Authors’ contributions

Conceptualization, GO, IK, NB and OF; Formal analysis, GO; Investigation, GO; Methodology, GO; Project administration, OF; Visualization, GO; Writing–original draft, GO; Writing–review and editing, GO, IK, NB and OF. All authors read and approved the final manuscript.

## Funding

This study was partially supported by the Grant No. 20-015-00123 from Russian Foundation for Basic Research and the Grant No. 075-15-2019-1789 from the Ministry of Science and Higher Education of the Russian Federation allocated to the Center for Precision Genome Editing and Genetic Technologies for Biomedicine.

## Competing interests

The authors declare that they have no conflict of interest relating to the conduct of this study or the publication of this manuscript.

## Notes

### Competing Interest Statement

The authors have declared no competing interest.

## References

[1] F. Wahid, A. Shehzad, T. Khan, Y.Y. Kim, MicroRNAs: synthesis, mechanism, function, and recent clinical trials, Biochimica et Biophysica Acta (BBA)-Molecular Cell Research. 1803 (2010) 1231–1243.

[2] A. Hata, Functions of microRNAs in cardiovascular biology and disease, Annual Review of Physiology. 75 (2013) 69–93.

[3] G. Condorelli, M.V. Latronico, E. Cavarretta, microRNAs in cardiovascular diseases: current knowledge and the road ahead, Journal of the American College of Cardiology. 63 (2014) 2177–2187.

[4] D. Quiat, E.N. Olson, MicroRNAs in cardiovascular disease: from pathogenesis to prevention and treatment, The Journal of Clinical Investigation. 123 (2013) 11–18.

[5] N. Baulina, G. Osmak, I. Kiselev, N. Matveeva, N. Kukava, R. Shakhnovich, O. Kulakova, O. Favorova, NGS-identified circulating miR-375 as a potential regulating component of myocardial infarction associated network, Journal of Molecular and Cellular Cardiology. 121 (2018) 173–179.

[6] V.N. Garikipati, S.K. Verma, D. Jolardarashi, Z. Cheng, J. Ibetti, M. Cimini, Y. Tang, M. Khan, Y. Yue, C. Benedict, Therapeutic inhibition of miR-375 attenuates post-myocardial infarction inflammatory response and left ventricular dysfunction via PDK-1-AKT signalling axis, Cardiovascular Research. 113 (2017) 938–949.

[7] M. Uhlén, L. Fagerberg, B.M. Hallström, C. Lindskog, P. Oksvold, A. Mardinoglu, \AAsa Sivertsson, C. Kampf, E. Sjöstedt, A. Asplund, Tissue-based map of the human proteome, Science. 347 (2015).

[8] C.-H. Chou, S. Shrestha, C.-D. Yang, N.-W. Chang, Y.-L. Lin, K.-W. Liao, W.-C. Huang, T.-H. Sun, S.-J. Tu, W.-H. Lee, miRTarBase update 2018: a resource for experimentally validated microRNA-target interactions, Nucleic Acids Research. 46 (2017) D296–D302.

[9] D. Szklarczyk, J.H. Morris, H. Cook, M. Kuhn, S. Wyder, M. Simonovic, A. Santos, N.T. Doncheva, A. Roth, P. Bork, L.J. Jensen, C. von Mering, The STRING database in 2017: quality-controlled protein–protein association networks, made broadly accessible, Nucleic Acids Res. 45 (2017) D362–D368. https://doi.org/10.1093/nar/gkw937.

[10] A. Hagberg, P. Swart, D. S Chult, Exploring network structure, dynamics, and function using NetworkX, in: Pasadena. USA., 2008: p. 11.

[11] N. Peláez, R.W. Carthew, Biological robustness and the role of microRNAs: a network perspective, in: Current Topics in Developmental Biology, Elsevier, 2012: pp. 237–255.

[12] P. Shannon, A. Markiel, O. Ozier, N.S. Baliga, J.T. Wang, D. Ramage, N. Amin, B. Schwikowski, T. Ideker, Cytoscape: a software environment for integrated models of biomolecular interaction networks, Genome Research. 13 (2003) 2498–2504.

[13] B. Jassal, L. Matthews, G. Viteri, C. Gong, P. Lorente, A. Fabregat, K. Sidiropoulos, J. Cook, M. Gillespie, R. Haw, The reactome pathway knowledgebase, Nucleic Acids Research. 48 (2020) D498–D503.

[14] Y.-A. Chen, L.P. Tripathi, B.H. Dessailly, J. Nyström-Persson, S. Ahmad, K. Mizuguchi, Integrated pathway clusters with coherent biological themes for target prioritisation, PloS One. 9 (2014) e99030.

[15] F. Beier, R.F. Loeser, Biology and pathology of Rho GTPase, PI-3 kinase-Akt, and MAP kinase signaling pathways in chondrocytes, J Cell Biochem. 110 (2010) 573–580. https://doi.org/10.1002/jcb.22604.

[16] E. Gutierrez, I. Cahatol, C.A.R. Bailey, A. Lafargue, N. Zhang, Y. Song, H. Tian, Y. Zhang, R. Chan, K. Gu, A.C.C. Zhang, J. Tang, C. Liu, N. Connis, P. Dennis, C. Zhang, Regulation of RhoB Gene Expression during Tumorigenesis and Aging Process and Its Potential Applications in These Processes, Cancers (Basel). 11 (2019). https://doi.org/10.3390/cancers11060818.

[17] T. Nakamura, M. Komiya, K. Sone, E. Hirose, N. Gotoh, H. Morii, Y. Ohta, N. Mori, Grit, a GTPase-activating protein for the Rho family, regulates neurite extension through association with the TrkA receptor and N-Shc and CrkL/Crk adapter molecules, Mol Cell Biol. 22 (2002) 8721–8734. https://doi.org/10.1128/mcb.22.24.8721-8734.2002.

[18] A. Murakami, M. Maekawa, K. Kawai, J. Nakayama, N. Araki, K. Semba, T. Taguchi, Y. Kamei, Y. Takada, S. Higashiyama, Cullin-3/KCTD10 E3 complex is essential for Rac1 activation through RhoB degradation in human epidermal growth factor receptor 2-positive breast cancer cells, Cancer Sci. 110 (2019) 650–661. https://doi.org/10.1111/cas.13899.

[19] S. Zhu, H. Liu, Y. Wu, B.C. Heng, P. Chen, H. Liu, H.W. Ouyang, Wnt and Rho GTPase signaling in osteoarthritis development and intervention: implications for diagnosis and therapy, Arthritis Res Ther. 15 (2013) 217. https://doi.org/10.1186/ar4240.

[20] A.B. Jaffe, A. Hall, Rho GTPases: biochemistry and biology, Annu Rev Cell Dev Biol. 21 (2005) 247–269. https://doi.org/10.1146/annurev.cellbio.21.020604.150721.

[21] P. Aspenström, A. Fransson, J. Saras, Rho GTPases have diverse effects on the organization of the actin filament system, Biochem J. 377 (2004) 327–337. https://doi.org/10.1042/BJ20031041.

[22] G.A. Murphy, P.A. Solski, S.A. Jillian, P. Pérez de la Ossa, P. D’Eustachio, C.J. Der, M.G. Rush, Cellular functions of TC10, a Rho family GTPase: regulation of morphology, signal transduction and cell growth, Oncogene. 18 (1999) 3831–3845. https://doi.org/10.1038/sj.onc.1202758.

[23] A. Flentje, R. Kalsi, T.S. Monahan, Small GTPases and Their Role in Vascular Disease, Int J Mol Sci. 20 (2019). https://doi.org/10.3390/ijms20040917.

[24] L.-Y. Dong, X.-X. Qiu, Y. Zhuang, S. Xue, Y-27632, a Rho-kinase inhibitor, attenuates myocardial ischemia-reperfusion injury in rats, Int J Mol Med. 43 (2019) 1911–1919. https://doi.org/10.3892/ijmm.2019.4097.

[25] L.-L. Gong, L.-H. Fang, S.-B. Wang, J.-L. Sun, H.-L. Qin, X.-X. Li, S.-B. Wang, G.-H. Du, Coptisine exert cardioprotective effect through anti-oxidative and inhibition of RhoA/Rho kinase pathway on isoproterenol-induced myocardial infarction in rats, Atherosclerosis. 222 (2012) 50–58. https://doi.org/10.1016/j.atherosclerosis.2012.01.046.

[26] X. Qian, M. Zhu, W. Qian, J. Song, Vitamin D attenuates myocardial ischemia-reperfusion injury by inhibiting inflammation via suppressing the RhoA/ROCK/NF-ĸB pathway, Biotechnol Appl Biochem. 66 (2019) 850–857. https://doi.org/10.1002/bab.1797.

[27] Y. Dai, J. Song, W. Li, T. Yang, X. Yue, X. Lin, X. Yang, W. Luo, J. Guo, X. Wang, S. Lai, K.C. Andrade, J. Chang, RhoE Fine-Tunes Inflammatory Response in Myocardial Infarction, Circulation. 139 (2019) 1185–1198. https://doi.org/10.1161/CIRCULATIONAHA.118.033700.

[28] P. Patel, M. Parikh, H. Shah, T. Gandhi, Inhibition of RhoA/Rho kinase by ibuprofen exerts cardioprotective effect on isoproterenol induced myocardial infarction in rats, Eur J Pharmacol. 791 (2016) 91–98. https://doi.org/10.1016/j.ejphar.2016.08.015.

[29] M.S. Ali Sheikh, Overexpression of miR-375 Protects Cardiomyocyte Injury following Hypoxic-Reoxygenation Injury, Oxid Med Cell Longev. 2020 (2020) 7164069. https://doi.org/10.1155/2020/7164069.

[30] S. Aznar, J.C. Lacal, Rho signals to cell growth and apoptosis, Cancer Lett. 165 (2001) 1–10. https://doi.org/10.1016/s0304-3835(01)00412-8.

[31] G.-D. Hu, Y.-H. Chen, L. Zhang, W.-C. Tong, Y.-X. Cheng, Y.-L. Luo, S.-X. Cai, L. Zhang, The generation of the endothelial specific cdc42-deficient mice and the effect of cdc42 deletion on the angiogenesis and embryonic development, Chin Med J (Engl). 124 (2011) 4155–4159.

